# Spectroscopy and Machine Learning Based Rapid Point-of-Care Assessment of Core Needle Cancer Biopsies

**DOI:** 10.1101/745158

**Authors:** Krishna Nand Keshavamurthy, Dmitry V. Dylov, Siavash Yazdanfar, Dharam Patel, Tarik Silk, Mikhail Silk, Frederick Jacques, Elena N Petre, Mithat Gonen, Natasha Rekhtman, Victor Ostroverkhov, Howard I. Scher, Stephen B. Solomon, Jeremy C. Durack

## Abstract

Solid tumor needle biopsies are essential to confirm malignancy and assess for actionable characteristics or genetic alterations to guide treatment selection. Ensuring that sufficient and suitable material is acquired for tumor profiling, while minimizing patient risk, remains a critical unmet need. Here, we evaluated the performance characteristics of transmission optical spectroscopy for rapid identification of malignant tissue in core needle biopsies (CNB). Human kidney biopsy specimens (545 CNB from 102 patients, 5583 spectra for analysis) were analyzed directly on core biopsy needles with a custom-built optical spectroscopy instrument. Machine learning classifiers were trained to differentiate malignant from normal tissue spectra. Classifiers were compared using receiver operating characteristics analysis and sensitivity and specificity were calculated relative to a histopathologic gold standard. The best performing algorithm was the random forest (sensitivity 96% and 93%, specificity 90% and 93% at the level of individual spectra and full CNB, respectively). *Ex-vivo* spectroscopy paired with machine learning paves the way towards rapid and accurate characterization of CNB at the time of tissue acquisition and improving tumor biopsy quality.

## Introduction

The rate of molecular diagnostic discoveries has not only increased the number of solid tumor biopsies performed but also magnified the importance of these specimens for classifying cancer and guiding treatment selection to optimize patient outcomes.^1, 2^ Unfortunately, image-guided needle biopsies often fail to provide adequate material for complete characterization by cell morphology, staining properties, immunohistochemistry, or genetic signatures.^3^ Rapid evaluation of specimen quality and tumor yield at the time of a biopsy could provide critical feedback to the operator, leading to more accurate and efficient tumor assessment.

Immediate biopsy specimen evaluation using cytologic imprints (touch preparations) under light microscopy can improve sampling accuracy.^4, 5^ However, the resources and expertise necessary to provide rapid on-site sample assessment are far from ubiquitous.^6–8^ Glass slide touch-preparation techniques can also be deleterious to downstream processing by substantially depleting the CNB of neoplastic cells or disrupting tissue architecture.^9^ In pursuing cancer biopsy quality improvement, a variety of technologies and techniques have been considered for point-of-care *in vivo* or *ex vivo* tissue characterization.^10^ These include optical spectroscopy,^11^ x-ray imaging,^12^ confocal microscopy,^13–15^ structured illumination microscopy,^16^ Fourier transform infrared imaging,^17^ fluorescence microscopy,^18–21^ contrast-enhanced micrography,^22^ as well as diffuse reflectance, electrical impedance, and Raman spectroscopy.^23–26^ Use of these technologies in a clinical setting has been hampered by factors such as lengthy analytic times, tissue degradation, expense, on-site tissue staining, requirement for interpretive expertise, and challenges to workflow integration. Additionally, many of these technologies analyze tissues *in vivo*, which may be helpful for determining intraoperative surgical margins but is less informative for *ex vivo* tissue acquired from a needle biopsy.

To facilitate *ex vivo* CNB quality assessment, we developed a transmission optical spectroscopy instrument that leverages machine learning methods to rapidly characterize CNB samples. Designed for bedside or procedure room clinical workflow integration, this hardware and software platform can acquire spectra from CNBs that remain intact on the biopsy needle sampling trough in less than one minute. In this study, we trained machine learning algorithms to determine whether individual transmission spectra obtained at regular intervals along a biopsy sample belong to either tumor or benign tissue classes. The robustness of classification to confounding sample types including fat, blood, and necrotic tissue was analyzed using multi-class classifiers. Additionally, a full CNB classifier, developed from the individual spectral classifier, was used to differentiate tumor-containing from benign tissue biopsies. Histopathologic diagnoses provided by our institutional clinical pathology laboratory formed the ground truth for training all algorithms.

Using the above approach, relevant direct-to-operator feedback included analyzable specimen length, a magnified image of the tissue core, geometric proportion of the sample that contained malignant cells, and classification of the CNB sample as malignant or benign. Based on this feedback, an operator may choose to either make additional needle passes or conclude the biopsy procedure. After spectroscopic analysis, full CNB samples can then undergo standard of care diagnostic and molecular analysis, thus minimizing sources of pre-analytical variation such as percentage of samples containing low volume or no tumor.

## Material and Methods

### Biopsy protocol

This study was approved by the institutional review board and human biospecimen utilization committee. Surgically excised human kidney specimens from April 2013 through October 2014 were biopsied *ex vivo* in a tissue procurement facility immediately after partial or complete nephrectomy for renal cell carcinoma. The distinction between normal and tumor regions was made by gross examination of pathology specimens. Biopsies were obtained using 18-gauge side-notch, spring-loaded core needle devices with 20 mm-long sampling troughs exposed (Temno Evolution or Adjustable Coaxial Temno, CareFusion Corporation, San Diego, CA). Kidney tumor and normal renal parenchymal were sampled in addition to renal sinus fat, visibly necrotic tissues, and blood. The histopathologic diagnosis of each tumor was recorded for association with spectroscopy data.

### Optical spectroscopy

Biopsy samples were analyzed using a custom-built automated transmission optical spectrometer (Figure 1A). A deuterium-tungsten light source (Ocean Optics, DH-2000-DUV, Dunedin, FL) delivered broadband illumination in the ultraviolet-visible-near infrared (UV-VIS-NIR) range (200–1100 nm). Light passed through a correction filter (Newport, FSR-KG2, Irvine, CA) for uniform output across the lamp’s spectrum. To maximize transmission of UV signal, the illumination light was guided via a 600 μm core multi-mode solarization-resistant fiber (Ocean Optics, QP600-1-SR) and focused on the sample using a 10 mm focal length fused silica lens (Ocean Optics, 74-UV). Illumination intensity was controlled by neutral density filters (Thorlabs, NDUV series, Newton, NJ) selected from an addressable motorized filter wheel (Thorlabs, FW102C).

**Figure 1.**
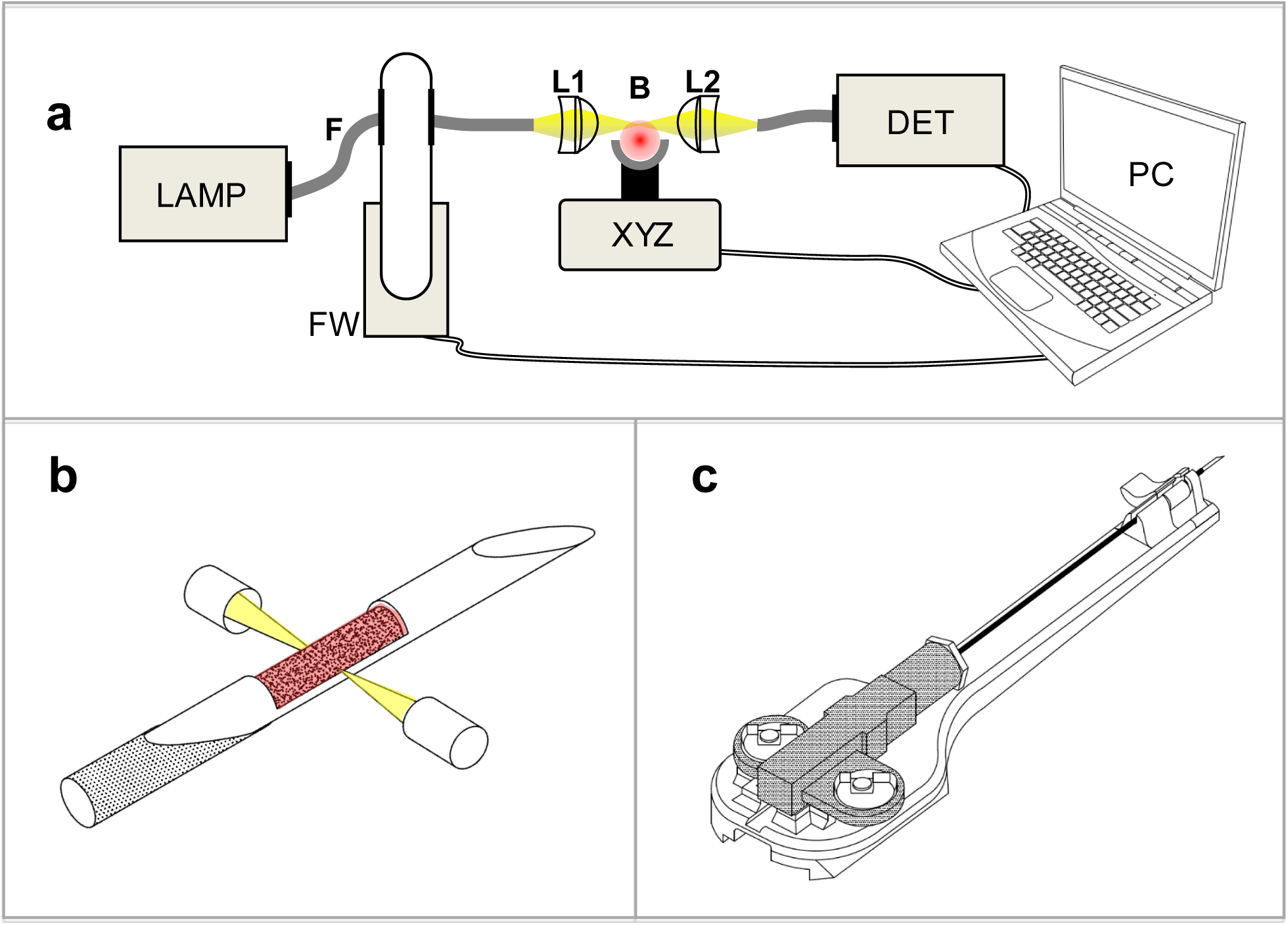
Schematic diagram of core biopsy spectroscopic imaging platform and subcomponents. Panel a) Transmission spectroscopy systems. Deuterium/tungsten light source (LAMP), Optical fiber (F), Filter wheel (FW), Lenses (L1, L2), Biopsy tissue core (B), Motorized translation stage (XYZ), Detector (DET), Laptop computer (PC). Panel b) Magnified and isolated schematic of core biopsy needle tip, tissue capture trough and light path for specimen imaging. Panel c) Schematic of 3D printed needle holder used for alignment of the needle tip with the illumination beam.

Imaging was performed directly across the CNB sampling trough (Figure 1B). A custom snap-in acrylonitrile butadiene styrene 3D-printed needle holder was used to align the tip of the needle with the illumination beam prior to scanning (Figure 1C). Light transmitted through the tissue sample was collected by fused silica lens (74-UV) and fiber-optically guided to a spectrometer coupled with a multi-channel array detector (Horiba, VS70, spectral window: 190–1000 nm, Kyoto, Japan).

Prior to scanning each biopsy sample, a dark current and a reference spectrum of the lamp were sequentially acquired. The needle was initially passed through the illumination beam to automatically detect sample boundaries, at which point a full scan was performed along the length of the specimen. During each scan, up to 21 transmission spectra were collected over 15 mm, at approximately 0.75 mm sampling increments. Similar to routine clinical CNBs, discontinuous samples (fragments) were commonly acquired, resulting in fewer than 21 spectra per sample. High dynamic range spectra were constructed by recording multiple exposures (10, 100, and 1000 msec integration times) merged into a single spectrum optimized for signal-to-noise across the entire 190–1000 nm detector sensitivity range. Instrument control and data acquisition were performed using a laptop computer with LabVIEW software (National Instruments, Austin, TX).

### Spectrum pre-processing

The acquired transmittance spectra were normalized to values between 0 and 1 by subtracting the dark spectrum and dividing by the reference spectrum. Low signal spectra due to obstruction of the light path by the needle, or saturated spectra due to 100% transmittance in unfilled sampling trough regions, as well as spectra outside the normalized population geometrical mean interquartile range were automatically rejected as outliers. Representative examples of normalized transmittance from kidney tumor, normal kidney, unclassifiable, and outlier spectra are shown in Figure 2. Principal component analysis^27^ was performed and the top 25 spectral components with the highest variance were extracted for further analysis.

**Figure 2.**
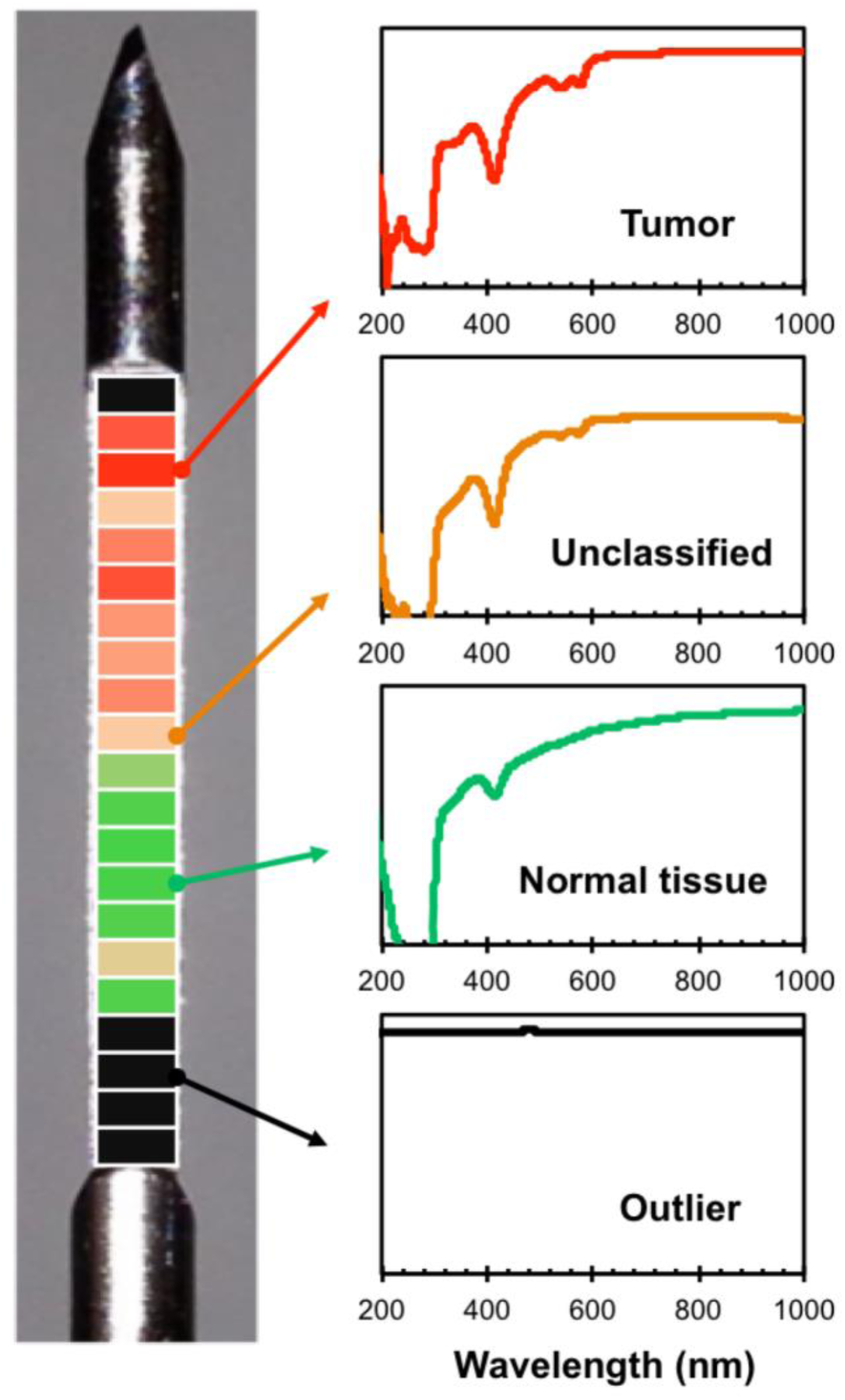
Example false-color tissue classification results displayed along the length of a biopsy sample obtained from the margin of a tumor and normal tissue. Normalized representative transmittance spectra are shown for tumor (red), unclassifiable (orange), normal tissue (green) and outlier (black) regions. The color scale indicates the probability (*p* = 0 to 1) that each imaging focal spot (horizontal band) represents tumor tissue. In this representative example, the spectral classification transitions from normal tissue (indicated by green, 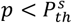) to tumor tissue (indicated by red, 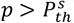). At the transition point, there is a spectrum with an unclassifiable value, 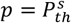 The proximal region of biopsy trough did not contain any tissue, resulting in outlier spectra (black) that were not classified.

### Automatic tissue classification using machine learning

All machine learning classifiers were trained on 80% of the data and tested on the remaining 20%. The training-test set split was randomized and stratified, and classifier hyperparameters were tuned using 10-fold stratified cross validation on the training set. All programming was carried out using MATLAB software.^28^

Machine learning classifiers were trained on the extracted principal spectral components to differentiate tumor spectra from normal tissue spectra. The ground truth labels for spectra were obtained from the histopathologic findings. The classification algorithms included LR, SVM, and RF.^25, 29^ The probability of a given spectrum belonging to tumor tissue was obtained as output from each classifier. The spectra were designated as either tumor or normal based on an optimal decision threshold on the probability of tumor 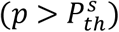. To test the robustness of classification to confounders, the analysis was repeated with three additional types of tissue spectra commonly obtained during clinical kidney mass CNBs: renal sinus fat, blood, and necrotic tumors. New multi-class classification models were trained to obtain probabilities of each spectrum belonging to one of the 5 classes. The multi-class output was further collapsed to a binary classification output, where a spectrum was designated as positive if classified as either viable or necrotic renal cell carcinoma and was designated as negative if classified as any of normal renal parenchyma, blood, or renal sinus fat.

Tissue classification was performed both at the individual spectrum and the full CNB sample levels. The probability of a full CNB sample representing predominantly tumor or normal tissue was computed by combining the likelihood of each of its constituent spectrum via a signal fusion approach and using naive Bayes assumption (Appendix A). The full CNB sample was designated as either tumor or normal based on an optimal decision threshold on the probability of tumor 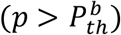

### Performance evaluation

Individual spectral and whole biopsy classifiers were quantitatively evaluated using ROC analysis and scalar performance metrics at their respective optimal decision thresholds. The optimal decision thresholds, 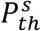 and 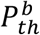, for the spectral and full CNB sample classifiers were computed using the Zweig and Campbell method^30^, where the slope of the optimal ROC point is computed based on a combination of mis-classification cost and prevalence analysis. The cost of misclassifying a tumor spectrum as normal tissue and vice-versa were assumed to be equal. McNemar’s statistical test was used to compare the performance of RF, LR, and SVM algorithms on spectral classification. Finally, to facilitate data visualization, false color “heat maps” for biopsy samples were generated by color coding the likelihood of constituent spectra belonging to tumor tissue. Visual representations indicating tumor presence and location within a CNB sample are displayed in Figures 2 and 6.

## Results

### Patients and Samples

Five hundred and forty-five CNBs were obtained from surgically resected kidneys from 102 patients undergoing partial or complete nephrectomy for renal cell carcinoma, resulting in 5583 spectra after pre-processing. Coring needles were advanced into selected regions based on direct tissue visualization and correlation with prior radiographic imaging studies. Sampled tissues included renal cell carcinoma, normal renal parenchyma, renal sinus fat, blood, and necrotic tumor (see Table 1 for the distribution of sampled tissues). Large tumor regions and normal (or non-tumor) tissue regions were sampled under the guidance of pathologists to ensure that samples contained either entirely tumor or entirely non-tumor tissues. The maximum scan duration was up to 60 sec for a full-trough biopsy sample.

**Table 1.** Distribution of biopsied tissue and total number of spectra obtained from each sample type.

| **Tissue** | **Specimens** | **Spectra** |
| --- | --- | --- |
| Renal cell carcinoma | 322 | 3176 |
| Normal renal parenchyma | 123 | 1618 |
| Renal sinus fat | 44 | 370 |
| Blood | 37 | 242 |
| Necrotic tumor | 19 | 177 |
| **Total** | **545** | **5583** |

### Classifier Performance

Figure 3 displays the classifier performance for binary (tumor vs. normal) and multi-class (confounders included) classification results. Of the three classification algorithms we evaluated (logistic regression (LR), support vector machine (SVM), and random forest (RF)), each performed well, though the random forest (RF) classifier achieved the highest area under the curve (AUC) of 0.98. The total computational time for classifying all the spectra from a biopsy sample and the whole biopsy sample itself was less than 1 sec.

**Figure 3.**
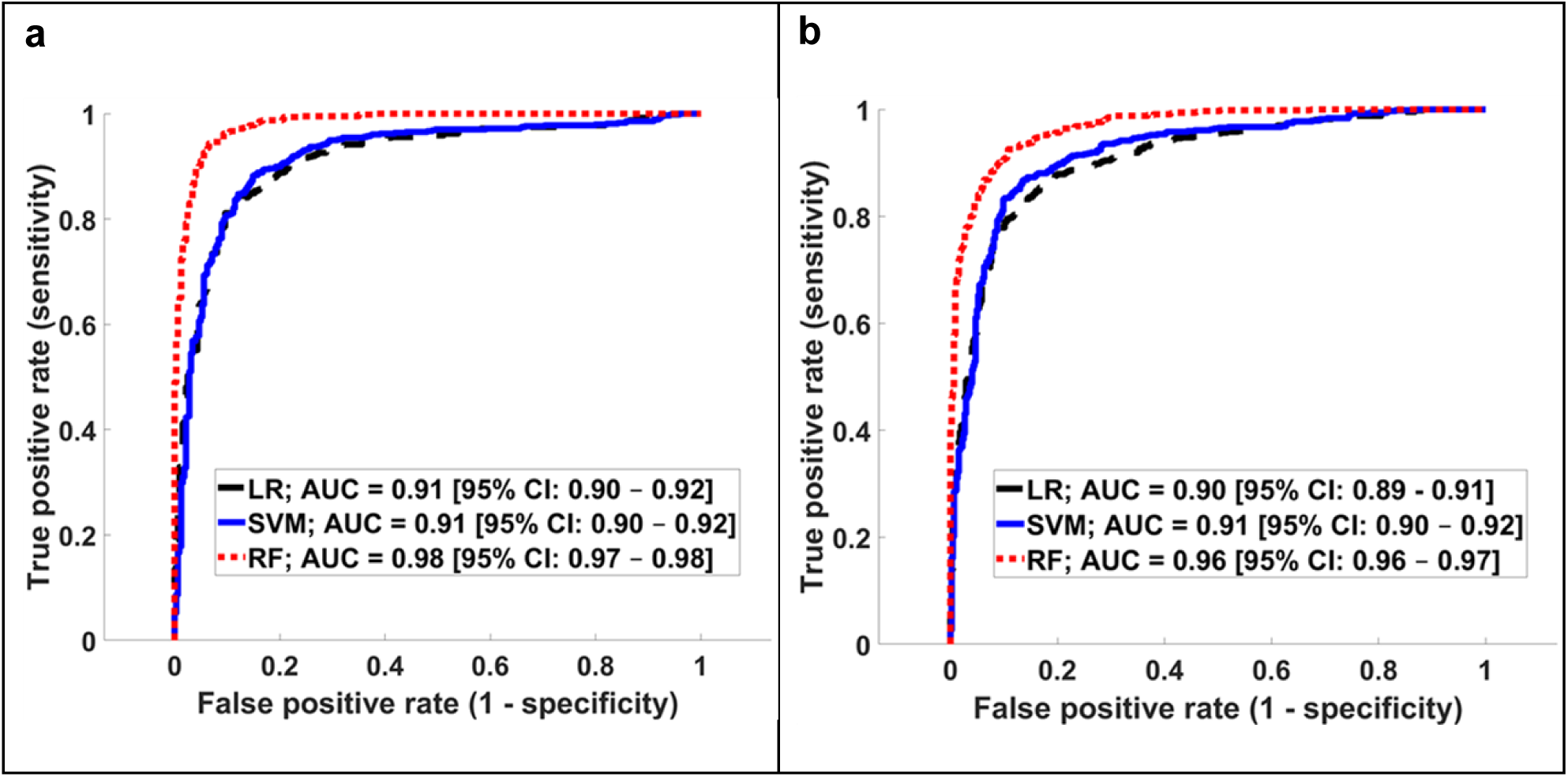
Receiver operating characteristic (ROC) analysis for spectral classification: a) tumor vs normal tissue; b) Multi-class classification of tumor and normal tissue with three additional confounding sample types: blood, fat, and necrotic tissue (right). The machine learning algorithms included logistic regression (LR), support vector machine (SVM), and random forest (RF).

### Robustness of Classification Analyzed using Multi-class Classifiers

Multi-class classification results were collapsed to a binary classification output, with tumor/necrotic tissues considered positive and normal tissues. Meanwhile, blood and fat considered negative. Here again, the RF achieved the best performance here as well with an AUC of 0.96.

Table 2 shows the scalar performance metrics (at optimal receiver operating curve (ROC) thresholds) for both the binary classification (top half) and the multi-class classification collapsed to a binary output (bottom half). RF achieved the best performance in both cases. At optimal ROC thresholds (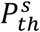 of 0.59 and 0.57), the sensitivities were 96% and 92% and specificities were 90% and 89% for the binary and multi-class analyses, respectively.

**Table 2.**
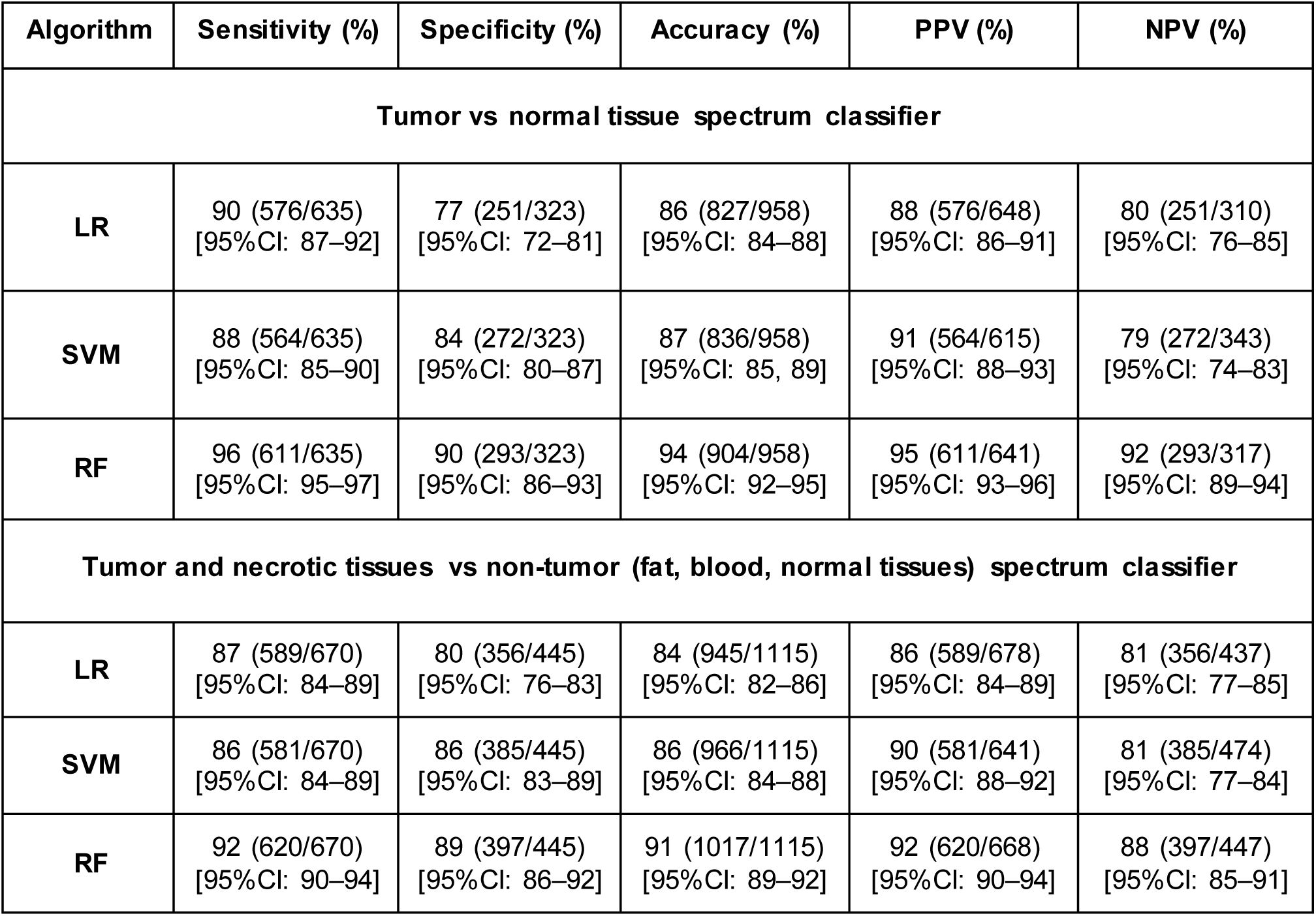
Statistical performance metrics comparing the spectral classifiers including logistic regression (LR), support vector machine (SVM), and random forest (RF). The top half of the table reflects binary tumor vs normal tissue spectral classification; the bottom half shows multiclass classification with confounders collapsed to a binary output (positive class: tumor, necrotic tissue spectra; negative class: fat, blood and normal tissue spectra). All the metrics were computed at their respective optimal ROC thresholds.

### Relative Performance of Classification Algorithms

Figure 4 shows the relative performance of the RF, LR, and SVM algorithms. Our null hypothesis that there was no difference in relative algorithm performance across all the compared metrics (error rate, sensitivity, and specificity) was examined. The difference between the performance of RF and the other algorithms was statistically significant in all the cases, and thus the null hypothesis was rejected. Taken together, our results demonstrated statistically significant superiority in terms of AUC, sensitivity, specificity, accuracy, PPV, and NPV for the RF algorithm over LR and SVM.

**Figure 4.**
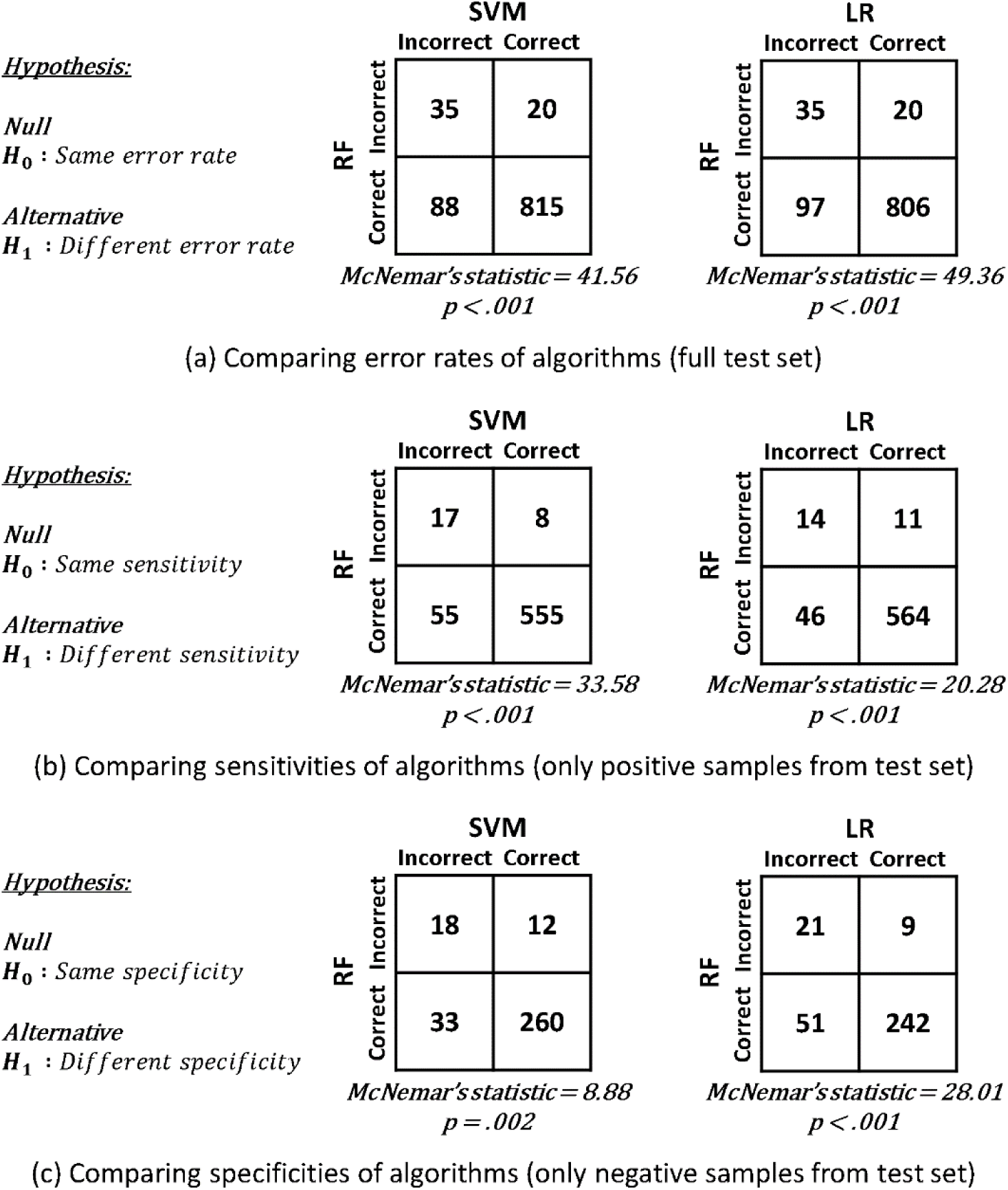
McNemar’s test for comparing random forest (RF) with support vector machine (SVM) and logistic regression (LR) spectral classifiers for each of the compared metrics: error rate (top), sensitivity (middle), and specificity (bottom), all computed at their respective optimal ROC thresholds on the held-out test set.

### Full CNB Classifier

Figure 5 shows the results of classifying the full CNB specimen, rather than single location individual spectra, as tumor or normal tissue. The best performing spectral classifier, RF, was used to build the full CNB classifier. At the optimal ROC threshold of 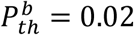, the sensitivity and specificity were 93% and 91% respectively. These results show that full CNBs can be classified with high fidelity.

**Figure 5.**
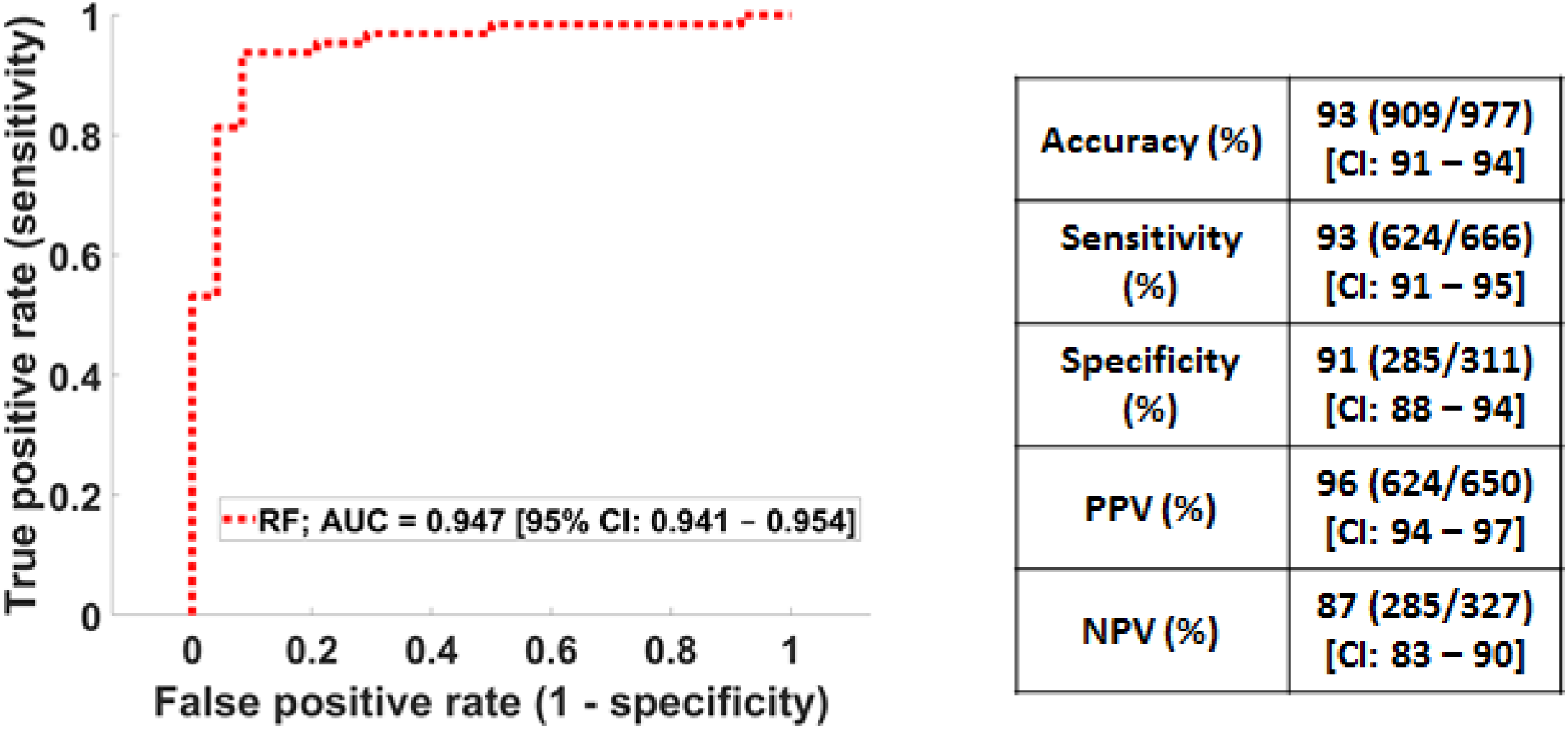
Whole biopsy classification using the random forest spectral classifier. The receiver operating characteristic curve (ROC) and the performance metrics at the optimal ROC threshold are shown.

**Figure 6.**
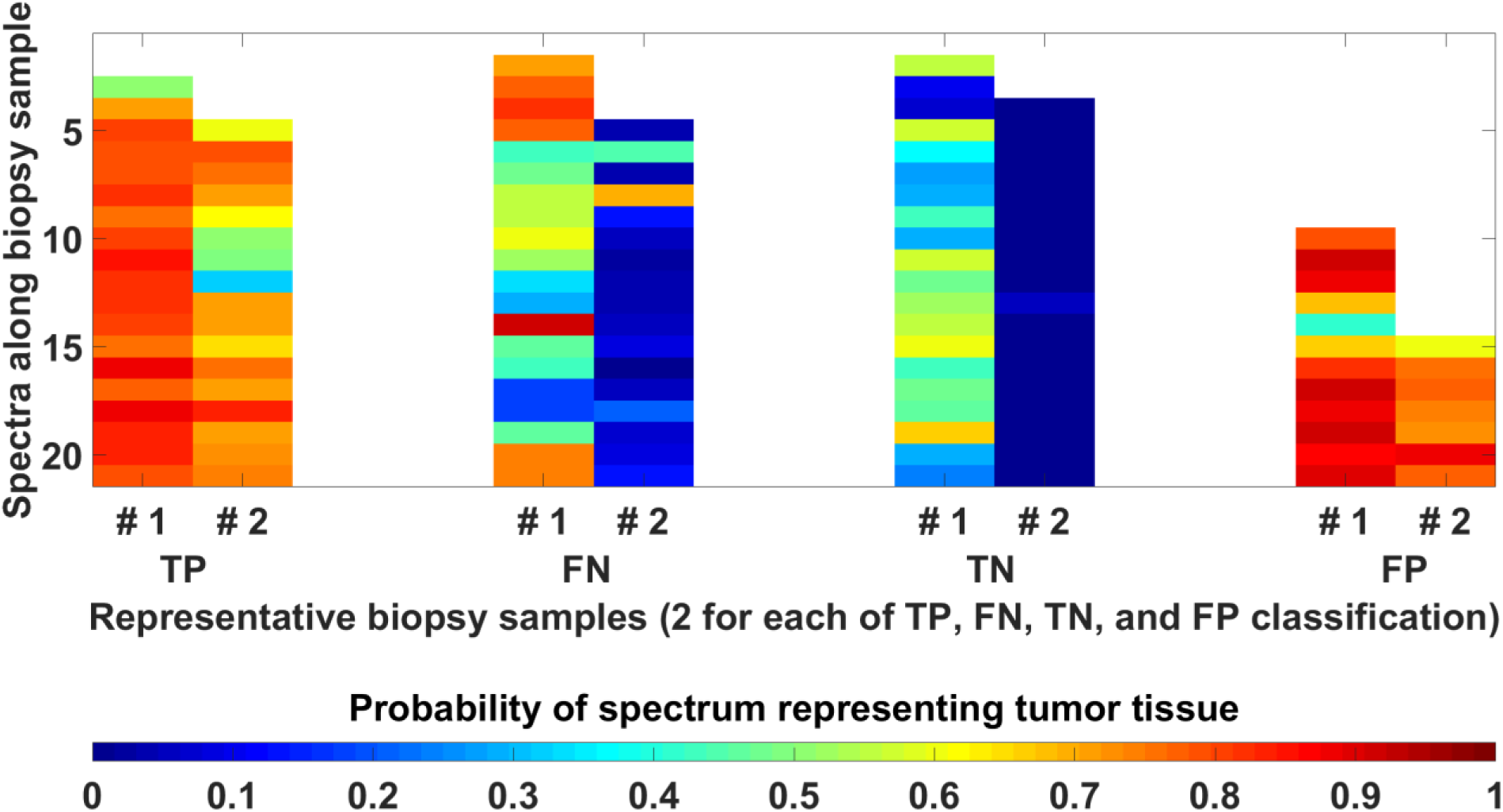
False-color heat maps of biopsy samples showing the likelihood and variability of constituent spectra belonging to tumor tissue representative of true positive (TP), false negative (FN), true negative (TN), and false positive (FP) classification.

### Heat Maps as Visualization Tools

Figure 6 shows false color heat maps correlating with the probability of tumor at each site of spectral acquisition along CNB samples, as determined by machine learning classification. Each CNB sample had up to 21 spectra depending upon the length of core tissue obtained from a needle pass and the number of spectra rejected as outliers. Two samples are presented in the figure for each of true positive (TP), false negative (FN), true negative (TN), and false positive (FP) cases based on individual spectra contributing to whole biopsy sample classification. TP and FP biopsy samples are largely green to red as their constituent spectra were predicted to belong to tumor tissue with high probability. Similarly, the TN and FN biopsy samples are largely blue to green as their constituent spectra were predicted to belong to tumor tissue with low probability. This visual representation reveals the presence and location of tumor and offers immediate visual feedback regarding tumor content.

## Discussion

Automated biological sample analyses that pair imaging with machine learning have shown to be both promising and challenging.^29^ Our study suggests that optical spectroscopy combined with machine learning can quickly and accurately characterize kidney CNB samples obtained for cancer diagnostics. The combination of an imaging modality, a reference library of tissue characteristics, a computational platform, and data networking may effectively close the physical and temporal gap between operating rooms or biopsy suites (tissue acquisition) and pathology departments (tissue analysis).

A variety of optical technologies for rapid high-resolution tissue imaging have been proposed and evaluated in the past, each with potential advantages and disadvantages in terms of portability, infrastructure requirements, ease of use, and cost.^10, 31, 32^ Relative to alternative systems that permit *in vivo* tissue characterization prior to biopsy using optical coherence tomography or elastography needle probes,^11^ the instrument studied here was designed to assess the *ex vivo* material obtained from the targeted lesion. This offers the advantage of providing feedback to the operator regarding the actual sample destined for downstream processing rather than a potential sampling target within the vicinity of the needle probe device. Utilizing this device, the extracted tissue is imaged while still present on the core biopsy needle trough immediately after it is acquired. Biopsy samples can be submitted to the pathology laboratory with no additional physical manipulation or risk of structural, cellular, or genetic damage. Additional samples can also be obtained from enriched tumor regions that may not have been collected otherwise.

Short scan durations and fast response times are important to minimize procedural delays, particularly when multiple biopsy samples are required for patients receiving sedation, or anesthesia. Spectroscopy scan time depends upon the number of focal spots for spectroscopy acquisition, which can be modified by the user if higher spatial resolution is desired. The present study data was acquired with scan times of less than 1 min per sample based on focal spots every 0.75 mm. Combined with the computational time for spectral and whole biopsy classification of less than 1 sec, our system is highly efficient relative to traditional imprint cytology slide preparation and review.

Direct feedback at the time of a biopsy can be leveraged to address challenges with standard of care biopsies today. Reducing undersampling could improve biopsy efficacy, while limiting oversampling could improve biopsy procedure safety. Rapid tumor mapping, combined with visualization tools such as the heat maps illustrated in this manuscript, could help the operator determine whether additional needle passes may be necessary. If indicated, supplemental biopsy material could then be obtained from a tumor-rich region or in a different region, depending upon the nature of the deficiency.

Our study was focused on the technical characterization of a spectroscopy instrument paired with automated classification. The potential for clinical translation and adoption of this or other similarly intended technologies is dependent upon a variety of factors. The “ASSURED” criteria (Affordable, Sensitive, Specific, User-friendly, Rapid and robust, Equipment-free, and Deliverable to end-users) are often cited to evaluate the potential of rapid point-of-care tests in resource or staff-limited environments.^10, 33^ Following these criteria, the technologies studied here are promising based on performance characteristics, speed, ease of use, and clear potential for integration into point-of-care environments. Nonetheless, robust clinical validation, true cost-effectiveness assessment, and data reproducibility studies will need to be performed. At present, the etiology of false positive and false negative sample classification is uncertain. This may be due to the relatively small size of the spectroscopic data and lack of availability of ground truth at the level of individual spectra, requiring ongoing contributions to increase specificity and sensitivity. Necrotic tumor tissues, inflammation, fibrosis, and residual blood on tissue samples also present a challenge for classification and are potential confounders. In particular, necrotic tumors may be considered either tumor positive or tumor negative, even by highly trained histopathologists depending upon whether small clusters of viable tumor cells are detected. Further studies in real-world biopsy settings will also be necessary to determine whether *ex vivo* tissue sample spectroscopic profiles may have been impacted by the time delay between surgical removal of the tumors and imaging. The high sensitivity and specificity achieved in this study suggests that tissue desiccation or ischemia does not significantly alter the spectroscopic signature of tumors; however, this requires additional assessment using freshly acquired specimens. While studies to date demonstrate high instrument sensitivity and specificity in kidney, the testing domain has been limited and requires expansion to a range of tumor types to generalize performance claims. To this end, a clinical trial utilizing this technology in a wide array of tissue and tumor types would be beneficial.

Lastly, a common criticism of machine learning classification algorithms to automate diagnoses is that there may be limited insight into the specific determinants used for classification. In this study, the underlying biological basis for critical spectral features is not known. Further efforts to improve classifier accountability may benefit from advances in the field of machine learning as a whole.

Point-of-acquisition tissue assessment using instruments such as the optical spectroscopy and machine learning platform described in this study could help to establish a new performance standard for rapid cancer biopsy assessment and automated needle biopsy quality control. The transformative potential of this technology is increasingly evident in the era of molecular oncology, where personalized cancer treatments require accurate and often repeated profiling of continually mutating cancer cell populations.

## Acknowledgments

We would like to acknowledge Joanne Chin, MFA, ELS, who provided editorial assistance with this manuscript, as well as Abhijit Patil, Rajesh Langoju, and Juan Pablo Cilia for their technical contributions. This work was supported in part by the NIH/NCI P30 Cancer Center Support Grant (P30 CA008748) and a Prostate Cancer Foundation Young Investigator research grant to J.C.D.

## Author Contributions

J.C.D., S.B.S., D.V.D., K.N.K.M., S.Y. conceived the research

K.N.K.M., S.Y., D.V.D., V.O. established the computational algorithms and processed the data

J.C.D., D.P., T.S., M.S., F.J. performed the tissue collection and spectroscopic imaging

M.G. provided expert statistical guidance

J.C.D., S.B.S. supervised the research

K.N.K.M., D.V.D., J.C.D., E.N.P. prepared the manuscript

All authors contributed to the manuscript.

K.N.K.M. and D.V.D. contributed equally

S.B.S. and J.C.D. jointly supervised this work

## Competing Interests

J.C.D., S.Y., V.O., D.V.D., S.B.S. Inclusion on patent applications related to spectroscopy instrument and data analytics.

J.C.D.: Scientific Advisory Board, Investor: Adient Medical; Chair, Society of Interventional Radiology Foundation

S.B.S.: Consultant: BTG, Johnson & Johnson, Adgero, XACT Robotics, Endoways, Aperture Medical

Grants: GE Healthcare, AngioDynamics, Elesta, Johnson & Johnson Shareholder: Johnson & Johnson, Aperture Medical, Innoblative, Immunomedics, Progenics

H.I.S.: Board of Directors, stock options: Asterias Biotherapeutics Consultancy/Advisory Board: Ambry Genetics Corporation, Konica Minolta,Inc., OncLive Insights, Physicians Education Resource, WCG Oncology (compensated); Amgen, ESSA Pharma Inc, Janssen Research & Development, LLC, Janssen Biotech, Inc. Menarini Silicon, Sanofi Aventis (uncompensated) Research Funding to the institution: Epic Sciences, Illumina, Inc, Innocrin Pharma, Janssen, Menarini Silicon, ThermoFisher Travel, Accommodations, Expenses: Amgen, Asterias Biotherapeutics, Clovis Oncology, ESSA Pharma Inc, Genome Profiling, LLC, Menarini Silicon, OncLive Insights, Physicians Education Resource, Prostate Cancer Foundation, Sanofi Aventis, WCG Oncology

V.O.: Employed by GE Global Research

## References

1. Marshall, D., Laberge, J.M., Firetag, B., Miller, T. & Kerlan, R.K. The Changing Face of Percutaneous Image-guided Biopsy: Molecular Profiling and Genomic Analysis in Current Practice. Journal of vascular and interventional radiology: JVIR (2013).

2. Ziv, E., Durack, J.C. & Solomon, S.B. The Importance of Biopsy in the Era of Molecular Medicine. Cancer J 22, 418–422 (2016).

3. Schneider, F. et al. Adequacy of core needle biopsy specimens and fine-needle aspirates for molecular testing of lung adenocarcinomas. Am J Clin Pathol 143, 193–200; quiz 306 (2015).

4. Tsou, M.H. et al. CT-guided needle biopsy: value of on-site cytopathologic evaluation of core specimen touch preparations. Journal of vascular and interventional radiology: JVIR 20, 71–76 (2009).

5. Kubik, M.J. et al. Diagnostic value and accuracy of imprint cytology evaluation during image-guided core needle biopsies: Review of our experience at a large academic center. Diagn Cytopathol (2015).

6. Balachandran, I. & Friedlander, M. Cytology workforce study: a report of current practices and trends in New York State. Am J Clin Pathol 136, 108–118 (2011).

7. Khurana, K.K. Telecytology and its evolving role in cytopathology. Diagn Cytopathol 40, 498–502 (2012).

8. Evans, A.J., Krupinski, E.A., Weinstein, R.S. & Pantanowitz, L. 2014 American Telemedicine Association clinical guidelines for telepathology: Another important step in support of increased adoption of telepathology for patient care. J Pathol Inform 6, 13 (2015).

9. Rekhtman, N. et al. Depletion of Core Needle Biopsy Cellularity and DNA Content as a Result of Vigorous Touch Preparations. Arch Pathol Lab Med 139, 907–912 (2015).

10. Boppart, S.A. & Richards-Kortum, R. Point-of-care and point-of-procedure optical imaging technologies for primary care and global health. Sci Transl Med 6, 253rv252 (2014).

11. Kennedy, K.M. et al. Needle optical coherence elastography for the measurement of microscale mechanical contrast deep within human breast tissues. J Biomed Opt 18, 121510 (2013).

12. Krupinski, E.A., Borders, M. & Fitzpatrick, K. Processing stereotactic breast biopsy specimens: impact of specimen radiography system on workflow. Breast J 19, 455–456 (2013).

13. Dobbs, J.L. et al. Feasibility of confocal fluorescence microscopy for real-time evaluation of neoplasia in fresh human breast tissue. J Biomed Opt 18, 106016 (2013).

14. Ragazzi, M. et al. Fluorescence confocal microscopy for pathologists. Mod Pathol 27, 460–471 (2014).

15. Dobbs, J.L. et al. Micro-anatomical quantitative optical imaging: toward automated assessment of breast tissues. Breast Cancer Res 17, 105 (2015).

16. Wang, M. et al. High-Resolution Rapid Diagnostic Imaging of Whole Prostate Biopsies Using Video-Rate Fluorescence Structured Illumination Microscopy. Cancer Res 75, 4032–4041 (2015).

17. Tiwari, S. et al. Towards Translation of Discrete Frequency Infrared Spectroscopic Imaging for Digital Histopathology of Clinical Biopsy Samples. Anal Chem 88, 10183–10190 (2016).

18. Mueller, J.L. et al. Quantitative Segmentation of Fluorescence Microscopy Images of Heterogeneous Tissue: Application to the Detection of Residual Disease in Tumor Margins. PLoS One 8, e66198 (2013).

19. Rivenson, Y. et al. Virtual histological staining of unlabelled tissue-autofluorescence images via deep learning. Nature biomedical engineering 3, 466–477 (2019).

20. Glaser, A.K. et al. Light-sheet microscopy for slide-free non-destructive pathology of large clinical specimens. Nature biomedical engineering 1 (2017).

21. Fereidouni, F. et al. Microscopy with ultraviolet surface excitation for rapid slide-free histology. Nature biomedical engineering 1, 957–966 (2017).

22. Im, H. et al. Design and clinical validation of a point-of-care device for the diagnosis of lymphoma via contrast-enhanced microholography and machine learning. Nature biomedical engineering 2, 666–674 (2018).

23. Spliethoff, J.W. et al. Real-time In Vivo Tissue Characterization with Diffuse Reflectance Spectroscopy during Transthoracic Lung Biopsy: A Clinical Feasibility Study. Clin Cancer Res 22, 357–365 (2016).

24. Shaikh, R. et al. A comparative evaluation of diffuse reflectance and Raman spectroscopy in the detection of cervical cancer. J Biophotonics 10, 242–252 (2017).

25. Huerta-Nunez, L.F.E. et al. A biosensor capable of identifying low quantities of breast cancer cells by electrical impedance spectroscopy. Scientific reports 9, 6419 (2019).

26. Orringer, D.A. et al. Rapid intraoperative histology of unprocessed surgical specimens via fibre-laser-based stimulated Raman scattering microscopy. Nature biomedical engineering 1 (2017).

27. Jolliffe, I.T. & Cadima, J. Principal component analysis: a review and recent developments. Philosophical transactions. Series A, Mathematical, physical, and engineering sciences 374, 20150202 (2016).

28. MathWorks MATLAB R2017a. (2014).

29. Patil, P.D., Hobbs, B. & Pennell, N.A. The promise and challenges of deep learning models for automated histopathologic classification and mutation prediction in lung cancer. J Thorac Dis 11, 369–372 (2019).

30. Zweig, M.H. & Campbell, G. Receiver-operating characteristic (ROC) plots: a fundamental evaluation tool in clinical medicine. Clinical chemistry 39, 561–577 (1993).

31. Taruttis, A. & Ntziachristos, V. Translational optical imaging. AJR Am J Roentgenol 199, 263–271 (2012).

32. Moriyama, E.H., Zheng, G. & Wilson, B.C. Optical molecular imaging: from single cell to patient. Clin Pharmacol Ther 84, 267–271 (2008).

33. Kettler, H., White, K. & Hawkes, S. Mapping the landscape of diagnostics for sexually transmitted infections: Key findings and recommendations. World Health Organization Special Programme for Research and Training in Tropical Diseases. Geneva, 2004.

