## Appendix A for "Spectroscopy and Machine Learning Based Rapid Point-of-Care Assessment of Core Needle Cancer Biopsies"

In this section, we present the derivation for computing the probability of the whole biopsy sample as belonging to one of tumor or normal tissue classes from its observed constituent spectra. We take a sensor fusion approach, where the spectra from the biopsy sample are treated as the output of different sensors. The likelihood of each sensor output (spectra) as belonging to one of tumor or normal class are fused/combined together using Naive Bayes assumption to compute the likelihood of the whole biopsy sample as belonging to tumor or normal class.

Let $B$ denote the binary random variable corresponding to the classification of the biopsy sample as belonging to one of tumor or normal tissue. i.e. let $B = \{T, N\}$, where $T$ stands for tumor tissue and $N$ stands for normal tissue; let $s_1, s_2, s_3 \dots s_n$ denote the observed constituent spectra from the biopsy sample, where $1 \leq n \leq 21$ is the number of observed constituent spectra from the biopsy sample (the maximum number of spectra from a biopsy sample in our experiments is 21); let $P(x)$ denote the probability of $x$. Then, our goal is to compute $P(B \mid s_1, s_2, s_3 \dots s_n)$. i.e. having observed the constituent spectra $s_1$, $s_2$, $s_3 \dots s_n$ from the biopsy sample, we want to compute the probability that the biopsy sample belongs to one of tumor or normal tissue.

Expanding the probability using Bayes' rule, we have

$$P(B \mid s_1, s_2, s_3 \dots s_n) = \frac{P(s_1, s_2, s_3 \dots s_n \mid B) * P(B)}{P(s_1, s_2, s_3 \dots s_n)}$$

$$P(B \mid s_1, s_2, s_3 \dots s_n) \propto P(s_1, s_2, s_3 \dots s_n \mid B) * P(B)$$

In this work, we assume that the spectra from a biopsy sample are independent given the class label. This is the Naive Bayes assumption [1]. Expanding out the probability further, we get

$$P(B \mid s_1, s_2, s_3 \dots s_n) \propto P(s_1 \mid B) * P(s_2 \mid B) * P(s_3 \mid B) \dots P(s_n \mid B) * P(B)$$

Using Bayes' rule on the conditional probabilities, we get

$$P(B \mid s_1, s_2, s_3 \dots s_n) \propto \frac{P(B \mid s_1) * P(s_1)}{P(B)} * \frac{P(B \mid s_2) * P(s_2)}{P(B)} \dots \frac{P(B \mid s_n) * P(s_n)}{P(B)} * P(B)$$

$$\propto \frac{P(B \mid s_1) * P(B \mid s_2) * P(B \mid s_3) \dots P(B \mid s_n)}{P(B)^{n-1}} * P(s_1) * P(s_2) * P(s_3) \dots P(s_n)$$

Now, for different values of $B = \{T, N\}$, the term $P(s_1) * P(s_2) * P(s_3) \dots P(s_n)$ is constant. Hence, we can simplify the proportionality further to get

$$P(B \mid s_1, s_2, s_3 \dots s_n) \propto \frac{P(B \mid s_1) * P(B \mid s_2) * P(B \mid s_3) \dots P(B \mid s_n)}{P(B)^{n-1}}$$

The left hand side of the above equation has the probability of the biopsy sample belonging to one of tumor or normal tissue, given its observed constituent spectra. On the right hand side, we see the product of the likelihoods from each of the spectra (the probability that the tissue is tumor or normal given the spectrum) in the numerator. This term combines the likelihood from different sensors (spectra). The denominator has the marginal probability of a biopsy sample being tumor or normal tissue, raised to the power of 1 less than the total number of spectra from the sample. In our experiments, the marginal probability of a biopsy sample being tumor or normal tissue is computed based on the ratio of the total number of tumor and normal biopsy samples in our data set. This completes the derivation.

References:

[1] Hand, D. J.; Yu, K. (2001). "Idiot's Bayes — not so stupid after all?". International Statistical Review. 69 (3): 385–399. doi:10.2307/1403452. ISSN 0306-7734. JSTOR 1403452.
